## Supplementary material for "Tiled-ClickSeq for targeted sequencing of complete coronavirus genomes with simultaneous capture of RNA recombination and minority variants": SData 2: FASTQ_R1_Split_from_primersBED_16Sep20.py

```
#!/bin/python3
import argparse
import gzip
parser = argparse.ArgumentParser()
parser.add_argument("R1Data", help="Root of Datafiles")
parser.add_argument("R2Data", help="Report Name")
parser.add_argument("BED", help="Report Name")
parser.add_argument("Output_Dir", help="Report Name")

#parser.add_argument("--Replicates", help="Report Name")
#parser.add_argument("--Conditions", help="Report Name")
#parser.add_argument("--Fraction", help="Report Name")
#parser.add_argument("--MinCount", help="Report Name")
args = parser.parse_args()

R1Data = str(args.R1Data)
R2Data = str(args.R2Data)
Primers = str(args.BED)
Output_Dir = str(args.Output_Dir)

def Rev_Comp(Seq):
        Seq = Seq.upper()
        basecomplement = {'A': 'T', 'C': 'G', 'G': 'C', 'T': 'A', 'N': 'N'}
        letters = list(Seq)
        letters = [basecomplement[base] for base in letters]
        return ''.join(letters)[::-1]

PrimerDict  = {}
AllPrimers = []
class Primer(object):
    ##Example = '''NC_045512.2     1876    1901    CoV_73  1       -       TACAACACGAGCAGCCTCTGAT'''
    def __init__(self, Ref, Start, Stop, Name, Score, Dir, Seq):
        self.Ref = Ref
        self.Name = Name
        self.Start = int(Start)
        self.Stop = int(Stop)
        self.Dir = Dir
        self.Seq = Seq
        self.Count = 0
        
    def __str__(self):
        return  str([self.Ref,
                self.Start,
                self.Stop,
                self.Dir,
                self.Seq])

#unknownprimer = open(Output_Dir + '/unknownprimer.split.sam', 'a')    
print("Reading in Primers...")
with open(Primers, 'r') as In:
    data = In.readline()
    while data:
        data = data.split()
        AllPrimers.append(data[6])
        PrimerDict[data[6]] = Primer(data[0], data[1], data[2], data[3], data[4], data[5], data[6])
        #str(data[3]) = open(Output_Dir + '/' + data[3] + '.split.sam', 'a')
        data = In.readline()

R2ReadDict = {}
class R2Read(object):
    def __init__(self, Name):
        self.Name = Name
        self.Seq = ''
        self.PrimerName = ''
        
    def __str__(self):
        return  str([self.Name, self.Seq])

m=0
n=0
p=0
print("Reading in R2 Data...")
with gzip.open(R2Data, 'rt') as In:
    Name = In.readline()
    while Name:
        Name = Name.split()[0]
        Seq = In.readline()
        In.readline()
        Quals = In.readline()
        R2ReadDict[Name] = R2Read(Name)
        Test = [i in Seq[:30] for i in AllPrimers]
        if any(Test):
            if sum(Test) == 1:
                ##Unique primer found
                R2ReadDict[Name].PrimerName = PrimerDict[AllPrimers[Test.index(True)]].Name
                R2ReadDict[Name].Seq = AllPrimers[Test.index(True)]
                #print(Name, Seq)
                n+=1
            else:
                R2ReadDict[Name].PrimerName = 'unknownprimer'
                print('Abiguous primer: ', Name, Seq)
                m+=1
        else:
            R2ReadDict[Name].PrimerName = 'unknownprimer'##write to unknown
            p+=1
        Name = In.readline()
        
print(str(m) + ' reads with ambiguous primer')
print(str(n) + ' reads with one primer')
print(str(p) + ' reads with no primer')

n=0
m = len(R2ReadDict)
n5 = m/20

Unknown = 0
print("Reading in and writing out R1 Data...")
with gzip.open(R1Data, 'rt') as In:
    Name = In.readline()
    while Name:
        n += 1
        if n % n5 == 0:
            print(" .... %s Percent of Reads Processed .... " % (n/n5))
        Name = Name.split()[0]
        Seq = In.readline()
        Third = In.readline()
        Quals = In.readline()
        try:
            Primer = R2ReadDict[Name].PrimerName + '.' + Rev_Comp(R2ReadDict[Name].Seq) + ''
            PrimerDict[R2ReadDict[Name].Seq].Count += 1
        except:
            Primer = 'unknownprimer.NNN.'
            Unknown += 1
        output = Output_Dir + '/' + Primer + '.fastq'
        with open(output, 'a') as Out:
            Out.write(Name + '\n' + Seq + Third + Quals)
        Name = In.readline()

##Make Report
with open(Output_Dir + '/log.txt','w') as Out:
    Out.write("Primer Name" + '\t' +
              "Sequence" + '\t' + 
              "Count in Dataset" + '\n')
    Out.write("Unknown Primer" + '\t' +
              "N" + '\t' + 
              str(Unknown) + '\n')
    for i in PrimerDict:
        Out.write(PrimerDict[i].Name + '\t' +
                  PrimerDict[i].Seq + '\t' + 
                  str(PrimerDict[i].Count) + '\n')
```
