## Supplementary material for "Tiled-ClickSeq for targeted sequencing of complete coronavirus genomes with simultaneous capture of RNA recombination and minority variants": SData 2: Mismatches_Qual-filter_Jan2021_fixed.py

```
import math
import re
import argparse
import numpy as np
import matplotlib.pyplot as plt
parser = argparse.ArgumentParser()
import gzip
parser.add_argument("Pileup", help="Description")
parser.add_argument("Output", help="Description")
parser.add_argument("Quals", help="Min PHRED score")
parser.add_argument("--MinCov", help= "Description")
args = parser.parse_args()

QualLim = int(args.Quals) + 33
if args.MinCov:
    MinCov = int(args.MinCov)
else:
    MinCov = 1
Out = open(str(args.Output), 'w')
Out.write("Ref\tPosition\tRefNuc\tCoverage\tA\tT\tG\tC\tErrorrate\t\n")

All_Data = []
All_Data.append(["Ref", "Coverage", "A", "T", "G", "C"])

Nucs = ['A','T','G','C','a','t','g','c']
Bases = ['.',',','A','T','G','C','N']
with gzip.open(str(args.Pileup), 'rt') as In:
        All_Data.append([0,0,0,0,0,0])
        Data = In.readline().split()
        while Data:
#          try:
            #Read Data
            Ref = Data[0]
            Position = int(Data[1])
            RefNuc = Data[2]
##            Coverage = float(Data[3])
            Seqs = Data[4].upper()
            Quals = Data[5]
            while len(All_Data) <= Position:
                    All_Data.append([0,0,0,0,0,0])
            Seqs_filt = []
            #remove read mapping quals
#            if '^' in Seqs:
#                    x,y = Seqs.split('^',1)
#                    y = y.split('^')
#                    y = [i[1:] for i in y]
#                    y = ''.join(y)
#                    Seqs = x+y
#            else:
#                    pass
            if '^' in Seqs:
                    x,y = Seqs.split('^',1)
                    y = y.split('^')
                    y = [i[1:] for i in y]
                    y = ''.join(y)
                    Seqs = x+y
            else:
                    pass
            InDels = ['-', '+']
            for j in InDels:
                    if j in Seqs:
                            Seqs,y = Seqs.split(j,1)
                            y = y.split(j)
                            for k in y:
                                Ks = re.split('(\d*)',k,1)
                                Seqs += Ks[2][int(Ks[1]):]
                    else:
                            pass
            Seqs = Seqs.replace('$','')
            #Seqs = Seqs.replace('^','')
            Seqs = Seqs.replace('~','')
#            Seqs = Seqs.replace('*','')
#            Seqs = Seqs.replace('>','')
#            Seqs = Seqs.replace('<','')
            if len(Seqs) != len(Quals):
                print(Position, Data)
#                break
#            for i in Seqs:
#                if i in Bases:
#                    Seqs_filt.append(i)
#            if len(Seqs) != len(Seqs_filt):
            n=0
            Seqs_filt = []
            for i in range(len(Seqs)):
                Qual = Quals[i]
                if ord(Qual) > QualLim:
                    if Seqs[i] != '>' and Seqs[i] != '<':
                        Seqs_filt += Seqs[i]
                    else:
                        pass
                else:
                    pass
            Coverage = float(len(Seqs_filt))
            All_Data[Position][0:2] = [RefNuc, Coverage]
            if RefNuc == "A":
                    for i in range(len(Seqs_filt)):
                            if Seqs_filt[i] != '.' and Seqs_filt[i] != ',':
                                         if Seqs_filt[i] == "T":
                                                All_Data[int(Position)][3] += 1
                                         elif Seqs_filt[i] == "G":
                                                All_Data[int(Position)][4] += 1
                                         elif Seqs_filt[i] == "C":
                                                All_Data[int(Position)][5] += 1
                                         else:
                                                 pass
                            else:
                                 pass
                            n+=1
            elif RefNuc == "T":
                        for i in range(len(Seqs_filt)):
                            #Qual = Quals[n]
                            if Seqs_filt[i] != '.' and Seqs_filt[i] != ',':
                                #if ord(Qual) > QualLim:
                                         if Seqs_filt[i] == "A":
                                                All_Data[int(Position)][2] += 1
                                         elif Seqs_filt[i] == "G":
                                                All_Data[int(Position)][4] += 1
                                         elif Seqs_filt[i] == "C":
                                                All_Data[int(Position)][5] += 1
                                         else:
                                                 pass
                                #else:
                                 #   pass
                            else:
                                 pass
                            n+=1
            elif RefNuc == 'G':
                  for i in range(len(Seqs_filt)):
                            #Qual = Quals[n]
                            if Seqs_filt[i] != '.' and Seqs_filt[i] != ',':
                                #if ord(Qual) > QualLim:
                                         if Seqs_filt[i] == "A":
                                                All_Data[int(Position)][2] += 1
                                         elif Seqs_filt[i] == "T":
                                                All_Data[int(Position)][3] += 1
                                         elif Seqs_filt[i] == "C":
                                                All_Data[int(Position)][5] += 1
                                         else:
                                                 pass
                                #else:
                                    #pass
                            else:
                                 pass
                            n+=1
            elif RefNuc == "C":
                        for i in range(len(Seqs_filt)):
                            #Qual = Quals[n]
                            if Seqs_filt[i] != '.' and Seqs_filt[i] != ',':
                                #if ord(Qual) > QualLim:
                                         if Seqs_filt[i] == "A":
                                                All_Data[int(Position)][2] += 1
                                         elif Seqs_filt[i] == "T":
                                                All_Data[int(Position)][3] += 1
                                         elif Seqs_filt[i] == "G":
                                                All_Data[int(Position)][4] += 1
                                         else:
                                                 pass
                                #else:
                                    #pass
                            else:
                                 pass
                            n+=1
            else:
                    pass

            Mismatches = 0
            for i in range(2,6):
                  Mismatches += All_Data[int(Position)][i]
            if Coverage >= MinCov:
                ErrorRate = Mismatches/Coverage
            else:
                ErrorRate = 0 #'NA'
            Out.write(str(Ref) + '\t'
              + str(Position) + '\t'
              + str(RefNuc) + "\t"
              + str(int(Coverage)) + "\t"
              + str(All_Data[int(Position)][2]) + "\t"
              + str(All_Data[int(Position)][3]) + "\t"
              + str(All_Data[int(Position)][4]) + "\t"
              + str(All_Data[int(Position)][5]) + "\t"
              + str(ErrorRate) + '\t\n')
            #Out.write(str(Ref) + '_' + str(Position-0) + '_' + str(All_Data[Position][0]) + "_" + str(All_Data[Position][1]) + "_" + str(ErrorRate) + '\n')
            Data = In.readline().split()

 #         except:
 #             print(Data)
 #             break
Out.close()

TransversionIndex = ['AT','AG','AC','TA','TG','TC','GA','GT','GC','CA','CT','CG']
CountsIndex = ['A','T','G','C']
Transversions = [0]*12
Counts = [0]*4

GT = {}
GA = {}
GC = {}
CA = {}
CT = {}
CG = {}
AT = {}
AG = {}
AC = {}
TA = {}
TG = {}
TC = {}
n=0
GTn = []
GAn = []
GCn = []
CAn = []
CTn = []
CGn = []
ATn = []
AGn = []
ACn = []
TAn = []
TGn = []
TCn = []
MinCount = 100
for i in All_Data:
    if 'Ref' not in i and i[0] != 0 and i[0] != 'N':
        Counts[CountsIndex.index(i[0].upper())] += int(i[1])
    else:
        pass
    if i[0] == "A" and int(i[1]) > MinCount:
        AT[n] = i[3]/float(i[1])
        u = math.log10(AT[n]) if AT[n] !=0 else AT[n]
        ATn.append(u)
        Transversions[0] += i[3]
        AG[n] = i[4]/float(i[1])
        u = math.log10(AG[n]) if AG[n] !=0 else AG[n]
        AGn.append(u)
        Transversions[1] += i[4]
        AC[n] = i[5]/float(i[1])
        u = math.log10(AC[n]) if AC[n] !=0 else AC[n]
        ACn.append(u)
        Transversions[2] += i[5]
    elif i[0] == "T" and int(i[1]) > MinCount:
        TA[n] = i[2]/float(i[1])
        u = math.log10(TA[n]) if TA[n] !=0 else TA[n]
        TAn.append(u)
        Transversions[3] += i[2]
        TG[n] = i[4]/float(i[1])
        u = math.log10(TG[n]) if TG[n] !=0 else TG[n]
        TGn.append(u)
        Transversions[4] += i[4]
        TC[n] = i[5]/float(i[1])
        u = math.log10(TC[n]) if TC[n] !=0 else TC[n]
        TCn.append(u)
        Transversions[5] += i[5]
    elif i[0] == "G" and int(i[1]) > MinCount:
        GA[n] = i[2]/float(i[1])
        u = math.log10(GA[n]) if GA[n] !=0 else GA[n]
        GAn.append(u)
        Transversions[6] += i[2]
        GT[n] = i[3]/float(i[1])
        u = math.log10(GT[n]) if GT[n] !=0 else GT[n]
        GTn.append(u)
        Transversions[7] += i[3]
        GC[n] = i[5]/float(i[1])
        u = math.log10(GC[n]) if GC[n] !=0 else GC[n]
        GCn.append(u)
        Transversions[8] += i[5]
    elif i[0] == "C" and int(i[1]) > MinCount:
        CA[n] = i[2]/float(i[1])
        u = math.log10(CA[n]) if CA[n] !=0 else CA[n]
        CAn.append(u)
        Transversions[9] += i[2]
        CT[n] = i[3]/float(i[1])
        u = math.log10(CT[n]) if CT[n] !=0 else CT[n]
        CTn.append(u)
        Transversions[10] += i[3]
        CG[n] = i[4]/float(i[1])
        u = math.log10(CG[n]) if CG[n] !=0 else CG[n]
        CGn.append(u)
        Transversions[11] += i[4]
    n+=1

Ts = np.array(Transversions).reshape(4,3)
Counts = np.array(Counts, dtype=float).reshape(4,1)
TransversionIndex = np.array(TransversionIndex).reshape(4,3)
#print(Ts/Counts * 10000)
#print(TransversionIndex)
def Average(Data):
    n = 0
    Total = 0
    for i in Data:
        if i < -1:
            Total += i
            n+=1
    return "%.2f" % (Total/float(n))

def roundup(x):
    return int(math.ceil(x / 100.0)) * 100

def MakeHist(Data, Name, bins, x,y,z):
    ax1 = fig.add_subplot(x,y,z)
    c = plt.hist(Data, bins)
    Max = roundup(max(c[0]))
    plt.text(-4.5, Max*0.8, Name)
    plt.text(-2, Max*0.8, Average(Data))
    ax1.set_xticks((-5,-4,-3,-2,-1,0))
    ax1.set_ylim(ymax = Max)

bins=50
plt.rcParams.update({'font.size': 6})

fig = plt.figure()
MakeHist(ATn, 'AT', bins, 4,3,1)
MakeHist(AGn, 'AG', bins, 4,3,2)
MakeHist(ACn, 'AC', bins, 4,3,3)

MakeHist(TAn, 'TA', bins, 4,3,4)
MakeHist(TGn, 'TG', bins, 4,3,5)
MakeHist(TCn, 'TC', bins, 4,3,6)

MakeHist(GAn, 'GA', bins, 4,3,7)
MakeHist(GTn, 'GT', bins, 4,3,8)
MakeHist(GCn, 'GC', bins, 4,3,9)

MakeHist(CAn, 'CA', bins, 4,3,10)
MakeHist(CTn, 'CT', bins, 4,3,11)
MakeHist(CGn, 'CG', bins, 4,3,12)

plt.savefig(str(args.Output) + '.pdf', format='pdf')
```
